# *Staphylococcus aureus* overcomes anaerobe-derived short-chain fatty acid stress via FadX and the CodY regulon

**DOI:** 10.1101/2022.02.21.481349

**Authors:** Joshua R. Fletcher, Alex R. Villareal, Mitchell Penningroth, Ryan C. Hunter

## Abstract

Chronic rhinosinusitis (CRS) is characterized by immune dysfunction, mucus hypersecretion, and persistent infection of the paranasal sinuses. While *Staphylococcus aureus* is a primary CRS pathogen, recent sequence-based surveys have found increased relative abundances of anaerobic bacteria, suggesting that *S. aureus* may experience altered metabolic landscapes in CRS relative to healthy airways. To test this possibility, we characterized the growth kinetics and transcriptome of *S. aureus* in supernatants of the abundant CRS anaerobe *Fusobacterium nucleatum*. While growth was initially delayed, *S. aureus* ultimately grew to similar levels as in the control medium. The transcriptome was significantly affected by *F. nucleatum* metabolites, with the *agr* quorum sensing system notably repressed. Conversely, expression of *fadX*, encoding a putative propionate coA-transferase, was significantly increased, leading to our hypothesis that short chain fatty acids (SCFAs) produced by *F. nucleatum* could mediate *S. aureus* growth behavior and gene expression. Supplementation with propionate and butyrate, but not acetate, recapitulated delayed growth phenotypes observed in *F. nucleatum* supernatants. A *fadX* mutant was found to be more sensitive than wild type to propionate, suggesting a role for FadX in the *S. aureus* SCFA stress response. Interestingly, spontaneous resistance to butyrate, but not propionate, was frequently observed. Whole genome sequencing and targeted mutagenesis identified *codY* mutants as resistant to butyrate inhibition. Together, these data show that *S. aureus* physiology is dependent on its co-colonizing microbiota and metabolites they exchange, and indicate that propionate and butyrate may act on different targets in *S. aureus* to suppress its growth.

**Importance:** *S. aureus* is an important CRS pathogen, yet is found in the upper airways of 30-50% of people without complications. The presence of strict and facultative anaerobic bacteria in CRS sinuses has recently spurred research into bacterial interactions and how they influence *S. aureus* physiology and pathogenesis. We show here that propionate and butyrate produced by one such CRS anaerobe, *F. nucleatum*, alter the growth and gene expression of *S. aureus*. We show that *fadX* is important for *S. aureus* to resist propionate stress, and that the CodY regulon mediates growth in inhibitory concentrations of butyrate. This work highlights the possible complexity of *S. aureus*-anaerobe interactions, and implicates membrane stress as a possible mechanism influencing *S. aureus* behavior in CRS sinuses.

## INTRODUCTION

Chronic rhinosinusitis (CRS) is an inflammatory condition of the sinuses that is broadly characterized by facial pain, mucus hypersecretion and accumulation, immune dysfunction, pathogen colonization, and persistent polymicrobial infection^1–6^. Although CRS affects up to 15% of the population and represents a substantial economic burden, its complexity has slowed development of new treatments and therapeutic strategies^7^. CRS patients are frequently prescribed antibiotics, yet many do not respond and require functional endoscopic sinus surgery (FESS) to remove accumulated mucus and inflamed mucosa that prevents proper sinus drainage^5^. Given the urgent threat of antimicrobial resistance among CRS microbiota, there is a critical need to better understand microbial community dynamics in the upper airways and how they may contribute to disease^8^.

*Staphylococcus aureus* is a frequently isolated CRS pathogen and is aggressively targeted by antibiotic therapy, yet, this bacterium is also prevalent and abundant in the upper airways of asymptomatic healthy individuals^9,10^. This seeming paradox suggests that colonization by *S. aureus* is not sufficient to drive disease, but rather that there may be important environmental cues in the upper airways that shift the lifestyle of *S. aureus* towards commensalism or pathogenesis. Indeed, in a genome-wide association study of *S. aureus* isolated from 28 CRS patients, few *S. aureus* genetic signatures were associated with CRS subtypes, suggesting that *S. aureus* pathogenesis in CRS is unlikely due to selection for increased production of a particular toxin^11^.

Application of culture-independent genomics to the study of CRS has led to a paradigm shift from a small number of etiologic bacterial species toward a polymicrobial basis of disease^4,5,12^. However, the role of the greater CRS microbiome in disease pathophysiology remains poorly understood. To address this knowledge gap, we recently surveyed 16S rRNA gene sequences in FESS-derived mucus from a cohort of CRS patients and found increased relative abundances of numerous anaerobic bacterial taxa, including many known to degrade mucin glycoproteins^6^. CRS bacterial communities enriched on mucins as a sole carbon source converged on similar profiles, typically dominated by a combination of *Streptococcus*, *Prevotella*, *Fusobacterium*, and *Veillonella*. Interestingly, *S. aureus* had a variety of growth phenotypes and gene expression patterns when cultured in supernatants from these enrichment communities, indicating that nutrient usage and metabolite release by co-colonizing microbiota can profoundly affect *S. aureus* physiology^6^. Enrichment supernatants that best supported *S. aureus* growth had low levels of short-chain fatty acids (acetate, propionate, butyrate; SCFAs) and undetectable levels of *Fusobacterium*, members of which are known for producing SCFAs as amino acid fermentation byproducts^13,14^. However, neither growth promotion nor inhibition could be ascribed to any one taxon or metabolite within these communities.

In this study, we extend our previous work by demonstrating that *F. nucleatum* metabolites impede *S. aureus* growth and repress transcription of the accessory gene regulator (*agr*) quorum sensing system while inducing a putative fatty acid degradation operon (*fadXEDBA*). We confirm that the SCFAs propionate and butyrate are sufficient to impair *S. aureus* growth and alter gene expression, while acetate had relatively little effect. We show that growth of a ∆*fadX* mutant is significantly attenuated in the presence of propionate only, despite differing from butyrate by only one carbon. Spontaneous resistance to growth inhibition by butyrate arose frequently, while we failed to obtain propionate resistant mutants. Genome sequencing of butyrate resistant mutants identified premature stop codons and in-frame deletions in the gene encoding the nutrient-responsive global regulator *codY*, indicating a connection between de-repression of the CodY regulon through nutrient limitation and SCFA resistance. These data suggest that certain anaerobes may influence CRS community structure by limiting *S. aureus* growth via propionate and butyrate production. In addition, they implicate the CodY regulon as a mechanism allowing *S. aureus* persistence in otherwise inhospitable anaerobic bacterial communities of the upper airways.

## MATERIALS AND METHODS

### Bacterial strains and growth conditions

Bacterial strains used throughout this study are shown in Table S1. Plasmids and primers used for mutagenesis and complementation can be found in Table S2 and S3, respectively. *Staphylococcus aureus* strains USA300 LAC, JE2 and *fadX*::tn transposon mutant (obtained from the Nebraska Transposon Mutant Library) were routinely cultured aerobically at 37ºC on LB agar (IBI Scientific IB49020) or with shaking at 220 rpm in LB broth, both supplemented as needed with 10 μg/mL chloramphenicol (Cm, Teknova C0325)^6,15^. *S. aureus* was also grown in cell free supernatants (CFS) of *Fusobacterium nucleatum* ATCC 25586 that had been cultured anaerobically for 48 h in BBL Brucella Broth (BD 2011088) supplemented with 250 μg/mL and 50 μg/mL of hemin and vitamin K (Hardy Diagnostics Z237), respectively. To test the effects of specific SCFAs on *S. aureus* growth and gene expression, the sodium salts of acetate (Fisher Scientific S209), propionate (Sigma P1880), or butyrate (Sigma 303410) were added at various concentrations to LB then passed through a 0.22 μm polyethersulfone (PES) filter prior to use. *Escherichia coli* strain One Shot TOP10 (ThermoFisher Scientific C404010) was used for cloning the *fadX* mutagenesis plasmid while *E. coli* DC10B was used for plasmid passaging to prevent cytosine methylation to facilitate easier transfer to *S. aureus*. *E. coli* strains were routinely grown on and in LB with 20 μg/mL Cm or 100 μg/mL ampicillin (Amresco 0339) as needed.

### Growth curves

Overnight cultures of wild type *S. aureus* and various mutants were diluted 1:100 in sterile PBS, then 5 μL was added to 195 μL of growth medium per well in a 96 well microtitre dish. Plates were incubated at 37ºC for 24 h in a BioTek Synergy H1 microplate reader for 24 h, with five seconds of orbital shaking performed prior to hourly OD_600_ readings.

### Biofilm quantification

The crystal violet staining of biofilm material was performed according to Merritt et al^16^. Briefly, overnight *S. aureus* LB cultures were centrifuged at 5,000 rpm for 5 minutes and washed once with sterile PBS. They were sub-cultured 1:100 into fresh media in a 96 well microtiter plate and incubated statically at 37ºC for 48 h, after which time the OD_600_ values were recorded in a BioTek Synergy H1 microplate reader. Plates were then inverted to remove the cultures, washed three times in water, and allowed to dry. The wells were stained with 0.1% w/v crystal violet for 15 minutes at room temperature. The crystal violet was removed, and the plates were washed a further three times in water and allowed to dry. The dye was solubilized with 30% acetic acid for 15 minutes and then the absorbance at 560 nm was recorded. The OD560 was normalized to the OD_600_ for each well to generate the final values.

### RNA extraction

For *S. aureus* growth in anaerobe cell-free supernatants and in LB with or without SCFA supplementation, 2 mL of growth medium was inoculated 1:100 with *S. aureus* overnight LB cultures and grown at 37ºC with shaking at 220 rpm. Growth of each culture was monitored until they reached an OD_600_ of ~0.2 to 0.3, after which they were centrifuged for 1 min at 14,000 rpm. Supernatants were discarded, and pellets suspended in 50 μL of fresh LB supplemented with 20 μg/mL of lysostaphin (Sigma-Aldrich L7386). These were then incubated in a 37ºC water bath for 15-20 min or until the suspension cleared (no longer than 30 min). One mL of TRIzol Reagent (ThermoFisher 15596018) was added to the lysate, pipetted gently until mixed, and incubated at room temperature (RT) for 5 min. 200 μL of chloroform (VWR 0757) was added per tube and samples were vigorously shaken for 15 s, then incubated at RT for 5 min. Phase separation was performed by centrifugation at 12,000 rpm for 15 min at 4ºC. ~500 μL of the aqueous phase was removed and added to 500 μL of 95% ethanol (Decon Laboratories, Inc. UN1170), vortexed for 5 s and incubated at RT for 5 min. RNA was then isolated using the Zymo RNA Clean & Concentrator-5 kit according to the manufacturer’s instructions, including an on-column DNase I treatment.

### NanoString analysis of *S. aureus* gene expression

A custom NanoString probe set (Table S4) was designed to target transcripts for several key *S. aureus* virulence factors, metabolic genes, and global regulators. The probe set also included six housekeeping genes for normalization. DNase I-treated RNA from *S. aureus* grown in triplicate to OD_600_ ~0.2 to 0.3 in control medium (Brucella Broth, BB) or 48 h cell-free supernatants from *F. nucleatum* (Fn CFS) was submitted to the University of Minnesota Genomics Center (UMGC) for hybridization to the custom probe set. Raw data were imported into the nSolver Advanced Analysis software package for normalization and differential gene expression analysis using default settings. Transcripts were considered differentially expressed if their levels changed by two-fold and the Benjamini-Hochberg adjusted p-value was less than 0.05. The heatmap was constructed using the pheatmap package (v.1.0.12) in R (v.4.1.0)^17^.

### Reverse transcription and quantitative real-time PCR

1.5 μg of RNA was reverse transcribed using M-MuLV Reverse Transcriptase (NEB M0253L) following the manufacturer’s protocol. cDNA was diluted 1:15 in sterile water prior to use in qRT-PCR using SsoAdvanced Universal SYBR Green Supermix (Bio Rad 1725271). PCR products for each gene being assayed (see Table S3 for primer sequences) were used to construct standard curves for quantification. To determine relative copy number, transcript levels were normalized to the housekeeping gene *gmk* (guanylate kinase), which was confirmed to be consistent across growth conditions.

### Construction of a *S. aureus* ∆*fadX* deletion mutant

~500 bp sequences flanking the *fadX* gene (*SAUSA300_0229*) were amplified by PCR using Q5 DNA Polymerase (NEB M0491L). For cloning purposes, the upstream amplicon contained a 5’ KpnI restriction site and the downstream amplicon contained a 3’ SacI site, while the internal ends contained complementary overhangs to facilitate overlap extension PCR to fuse the fragments together. The final product was a ~1 kb fragment encoding the first 11 codons and the stop codon of *fadX*. The amplicon was digested with KpnI-HF (NEB R3142S) and SacI-HF (NEB R3156S) and cloned into the temperature sensitive, counter-selectable mutagenesis plasmid pIMAY with T4 DNA ligase (NEB M0202)^18^. The ∆*fadX* plasmid was transformed into *E. coli* DC10B, then electroporated (2900 V, 25 μF, 100 Ω) in a 2 mm cuvette into *S. aureus.* The culture was grown at the plasmid replication permissive temperature of 28ºC with shaking for 4 h, after which time it was plated onto LB + 10 μg/mL Cm and incubated on the benchtop for 48 h. The single colony that was obtained was streaked onto two LB + Cm plates. One was incubated on the benchtop for 48 h to generate a freezer stock and the other was incubated at 37ºC. Cm-resistant colonies were screened via PCR for chromosomal integration of the plasmid. Positive colonies were grown in LB in the absence of selection at 37ºC for 24 h, subcultured 1:1000 into fresh LB for another 24 h, then plated onto LB + 1 μg/mL anhydrotetracycline hydrochloride (Sigma Aldrich 37919) for counter-selection via induction of the TetR-regulated *secY* antisense RNA and incubated overnight at 37ºC. Resultant colonies were patched onto fresh counter-selection agar and LB + Cm to screen for loss of the plasmid. Cm-sensitive clones were screened by PCR for loss of *fadX* coding sequence and four were confirmed as mutants by Sanger sequencing.

### HPLC method for extraction and measurement of organic acids in complex media

Reversed-phase high-performance liquid chromatography (HPLC) was used for the targeted quantification of acetate, butyrate, and propionate in cell free supernatants. Organic acids of interest were purified from complex media components through a modified liquid-liquid extraction method^19^. To account for analyte loss during extraction, 100μL of 0.2M succinate was added as an internal standard to 2mLs of each sample (9.5mM final concentration)^19^. After equilibration at room temperature for 5 min, 200μL of 12N HCl was added, and samples were vortexed for 15 seconds. 10mL of diethyl ether was then added to each sample before gently rolling them for a total of 30 min. After centrifugation for 5 min at 4000rpm, supernatants were transferred to a new extraction tube and 1mL of 1M NaOH was added before gently rolling for another 30 min. The resulting aqueous phase was extracted and transferred to an autosampler vial followed by addition of 100μL of 12N HCL before vortexing and storage at 4°C until analysis.

Samples were analyzed using a Dionex UltiMate 3000 UHPLC (Thermo Fisher) system equipped with a reversed-phase Acclaim Organic Acid (OA) Column (5μm, 120 A, 4.0 × 250mm). 8μL of each sample was injected and separation was achieved using a 32-minute isocratic instrument method (1.0 mL/min, 30°C) employing Na2SO4 (100mM, pH 2.6) as the mobile phase. The column was allowed to equilibrate for 8 min prior to sample injection. Absorbance was monitored at 210nm to identify compounds with a carboxylic acid functional group. Chromatograms were processed using Dionex Chromeleon 7 (Thermo Fisher) Chromatography Data System. Cobra Wizard was used to reproducibly identify and gate peaks of interest.

### Isolation and genome sequencing of butyrate resistant mutants

The ∆*fadX* mutant was grown for 24 hours in LB, serially diluted, plated onto LB supplemented with 200 mM sodium butyrate, and incubated overnight at 37ºC. Four large colonies were re-streaked onto the same media to confirm their ability to grow in the presence of butyrate. Genomic DNA was then isolated from our *S. aureus* USA300 LAC isolate and butyrate-resistant mutants using the PowerSoil Pro kit (Qiagen 47014). Overnight cultures were pelleted and suspended in 200 μL of an enzymatic lysis buffer (20 mM Tris-HCl, 2 mM EDTA, 1.2% (vol/vol) Triton X-100, and 20 μg/mL lysostaphin) for 30 min at 37ºC. Lysates were transferred to Power Bead tubes and the manufacturer’s protocol was followed with no further alterations. Genomic DNA was processed and sequenced on the Illumina MiSeq platform at the Microbial Genome Sequencing Center (MiGS, Pittsburgh, PA). Raw paired-end fastq files were imported into Geneious v.2022.0.1 and trimmed for quality using BBDuk with the following settings: Set ORDERED to true, k=27, mink=6, maskMiddle=true, hamming distance=1, right-ktrimming using 1 reference, quality-trimming both ends to Q30. Trimmed reads for our USA300 isolate were mapped to the *S. aureus* subsp. *aureus* USA300_FPR3757 reference genome (CP000255) using the Geneious Mapper on medium-low sensitivity with a minimum mapping quality of 20 and only mapping reads whose pair mapped appropriately nearby. The assembled USA300 genome was then used as the reference against which reads from the butyrate resistant mutants were mapped using the same quality trimming and mapping strategy detailed above. Putative SNPs and indels were detected using the “Find Variations/SNPS…” function within Geneious, requiring occurrence of the variation in >90% of reads, with a minimum of 10 reads.

### Data availability

Raw sequences were deposited at NCBI’s Sequence Read Archive (SRA) with the BioProject ID PRJNA798706.

## RESULTS

### *S. aureus* growth is impaired by *F. nucleatum*

In a previous study of upper airway (CRS) microbiota, we found that in patients (n=27) with detectable *Fusobacterium*, relative abundances of *Staphylococcus* were minimal or below the level of detection (Fig. 1A)^6^. When *S. aureus* was grown in supernatants derived from anaerobic enrichment cultures of CRS sinus mucus, it exhibited slower growth in those that had *Fusobacterium* as a core constituent genus of the enrichment community. These supernatants also contained higher levels of the short-chain fatty acids (SCFAs) propionate and butyrate^6^. Given these data, we hypothesized that *Fusobacterium* spp. exert an antagonistic effect on *S. aureus* through the production of SCFAs. To test this, we grew *S. aureus* USA300 LAC in filtered cell-free supernatants (CFS) from *F. nucleatum* ATCC 25586 grown for 48 h in Brucella Broth (BB) supplemented with hemin and vitamin K (Fig. 1B). *S. aureus* grew in Fn CFS, albeit slower than in the control medium (BB) with an increased lag phase. However, both cultures reached approximately similar OD_600_ values by 24 h, indicating that *S. aureus* was able to obtain sufficient nutrients over the course of the experiment. We reasoned that the extended lag phase was likely due to *F. nucleatum*-mediated depletion of an easily metabolizable nutrient source and/or the presence of an inhibitory metabolite(s) that *S. aureus* was able to adapt to over time. We tested the latter by measuring acetate, propionate, and butyrate content of the CFS before and after *S. aureus* growth (Fig. 1C). All three SCFAs were detected in *F. nucleatum* CFS (~5mM acetate, ~5mM propionate, ~15mM butyrate). After overnight growth (~16 h) in *F. nucleatum* supernatants, *S. aureus* cultures had increased acetate levels, while propionate and butyrate remained the same as in CFS alone, indicating that *S. aureus* does not actively metabolize the latter two SCFAs under these conditions. We interpret this to mean that *S. aureus* adapts to SCFAs by modifying its physiology rather than directly detoxifying them via degradation. The increased acetate levels are likely due to *S. aureus* utilization of glucose remaining in Fn CFS, as *F. nucleatum* preferentially ferments amino acids^20,21^. Given that *S. aureus* growth is similarly impaired when BB is supplemented with the sodium salts of acetate (5mM), propionate (5mM), and butyrate (15mM)(Fig. 1B), these data suggest that SCFAs may be key factors driving bacterial interactions in the CRS sinus environment, providing a mechanism by which *Fusobacterium* and other anaerobes may restrict *S. aureus* growth *in vivo*.

**Figure. 1.**
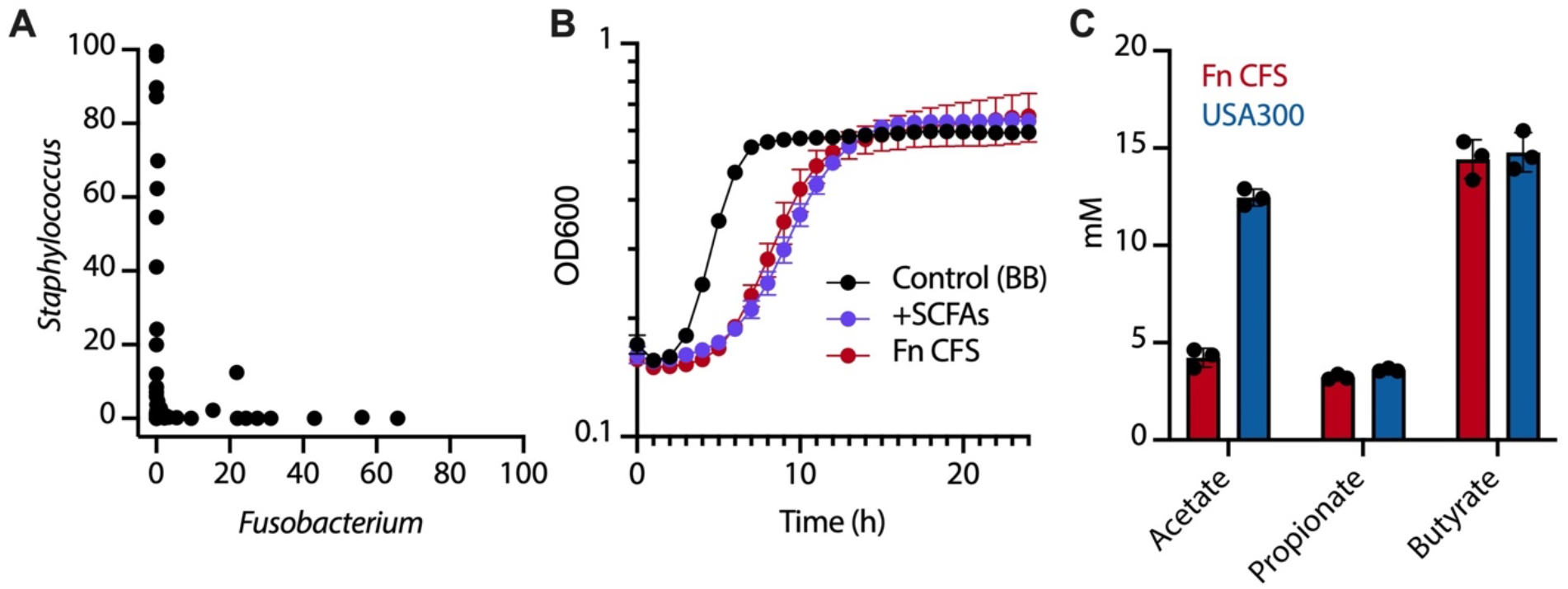
*S. aureus* growth is impaired in *F. nucleatum* supernatants. **A)** Relative abundances of *Fusobacterium* and *Staphylococcus* in sinus mucus from patients with chronic rhinosinusitis are inversely correlated (Lucas, et al 2021). **B)** Representative growth curve of *S. aureus* USA300 in brucella broth (BB, control), BB supplemented with 5mM acetate, 5mM propionate, and 15mM butyrate, and cell-free supernatants from *F. nucleatum* (Fn CFS). **C)** Production of SCFAs by *F. nucleatum* after 48h (Fn CFS) and their levels after *S. aureus* (USA300) growth in Fn CFS.

### *F. nucleatum* metabolites significantly alter *S. aureus* gene expression. We next determined how

*S. aureus* modified its transcriptome in *F. nucleatum* CFS. To do so, we performed a targeted analysis using a custom NanoString code set (Table S4) that included 34 genes encoding several known virulence factors, key metabolic genes, and master regulators of gene expression (Fig. 2A). Of these genes, we detected thirteen differentially expressed transcripts (≥2-fold change in expression and adjusted p-value<0.05); expression of *fadX*, *cidA*, *icaB*, and *gltB* increased while *nanA*, *alsS*, *lrgA*, *narG*, *agrA*, *hla*, *hld*, *saeR*, and *ldh1* decreased in *S. aureus* grown on CFS relative to BB alone (Fig. 2B). A number of other transcripts (*aur, fib, pgi, codY, opp3b,* and *arlR*) were statistically significant, but exhibited less than two-fold change in expression (Table S4). Decreased signaling through the quorum-sensing response regulator *agrA* in Fn CFS results in lower expression of the *hla* and *hld* genes, encoding alpha and delta hemolysins^22^. Neuraminate lyase, encoded by *nanA*, is induced by the presence of sialic acids and exhibited lower expression in Fn CFS, indicating that *F. nucleatum* likely utilized sialic acids present in Brucella Broth as a nutrient source^23,24^. Expression of the nutrient-sensing transcriptional regulator *codY* was reduced in Fn CFS by nearly 50% compared to the control medium, likely explaining the increase in the glutamate synthase subunit gene *gltB*^25^. We selected three genes (*fadX*, *agrA*, and *nanA*) for validation via quantitative reverse-transcription PCR and show that they were highly correlated with the NanoString results (Fig. S1). That nearly half of the transcripts in the NanoString probe set are differentially regulated in Fn CFS, including a number of major transcriptional regulators important for integrating metabolic cues and virulence gene expression, highlights the global nature of alterations to the *S. aureus* transcriptome. These data show that *S. aureus* physiology can be significantly influenced by the metabolic activity of a single anaerobic species and underscore the possible complexity of bacterial behaviors and interactions within a diverse CRS community.

**Figure. 2.**
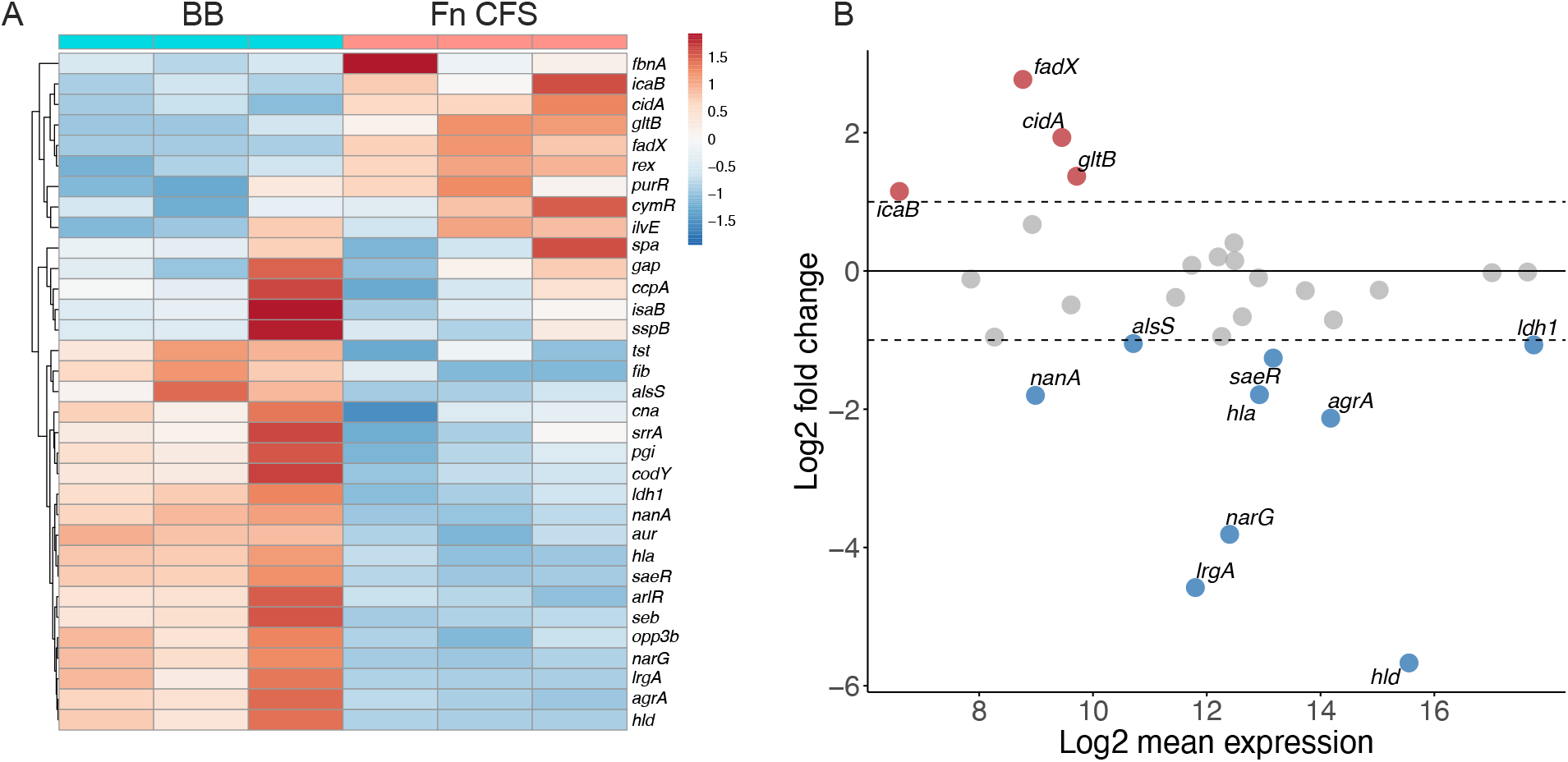
*F. nucleatum* metabolites significantly impact the *S. aureus* transcriptome. **A)** Heatmap depicting log10-transformed *S. aureus* gene expression in control media (BB) and *F. nucleatum* supernatant (CFS) as detected by NanoString. Genes were clustered with unsupervised hierarchical clustering. **B)** MA plot representation of *S. aureus* gene expression in Fn CFS relative to the control medium. Genes were considered significant if they had a log2 fold change ≥1 and a Benjamini-Hochberg adjusted p-value < 0.05.

### SCFAs significantly alter *S. aureus* gene expression

SCFAs, especially propionate and butyrate, have been reported to impair *S. aureus* growth and attenuate murine skin infections^26^. We therefore sought to determine if individual SCFAs were sufficient to drive some of the *S. aureus* gene expression patterns observed after growth in Fn CFS. To measure the effects of each SCFA on the *agr* quorum sensing system, we grew *S. aureus* carrying pAH1 (encoding P_*agr*_*-mCherry*) in LB or LB supplemented with acetate, propionate, or butyrate for 24 h and measured fluorescence intensity normalized to culture density (Fig. 3A). All three SCFAs led to decreased fluorescence, with propionate (p=0.0035) and butyrate (p<0.0001) significantly inhibiting reporter activity, while acetate (p=0.203) had the least effect. Given these observations, CRS bacterial communities dominated by *Fusobacterium* and other taxa that produce propionate and butyrate would not only be predicted to impede the growth of *S. aureus*, but also minimize the production of *agr*-regulated virulence factors.

**Fig. 3.**
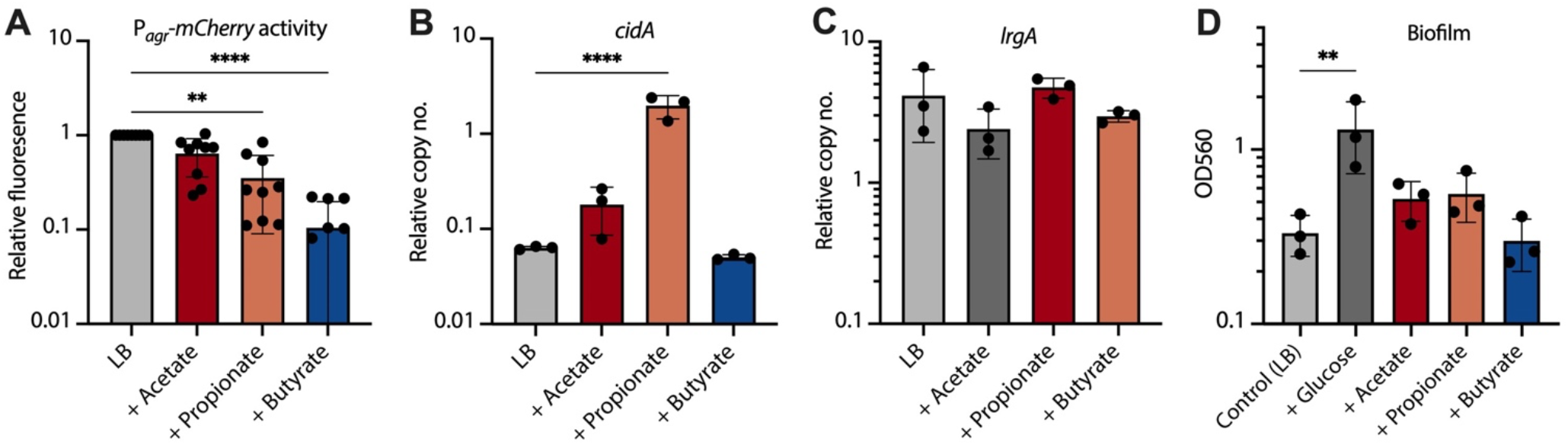
Propionate and butyrate repress the *S. aureus agr* system but fail to induce biofilm. **A)** *S. aureus* carrying pAH1 (P_*agr*_*-mCherry*) was grown for 24 hours in LB supplemented with 100 mM of sodium acetate, propionate, or butyrate (n=3 biological replicates with n=3 cultures per replicate). Fluorescence was measured and normalized to culture density for each replicate, then normalized to the LB controls. **B, C)** Expression of *cidA* and *lrgA* from *S. aureus* in LB supplemented with SCFAs (n=3). Copy number was determined via standard curve and normalized to the *gmk* housekeeping gene. **D)** Crystal violet assay quantifying biofilm formation in LB, LB supplemented with glucose (positive control for increased biofilm formation), or LB supplemented with 100 mM of each SCFA.

Given the reduction in *agr* quorum sensing activity and thus lack of repression of proteins involved in surface attachment, we hypothesized that SCFAs may be a pro-biofilm signal to *S. aureus*^27^. We performed qRT-PCR on *S. aureus* grown to OD_600_ ~0.2-0.3 in LB or LB supplemented with 100 mM of each individual SCFA to determine if biofilm-associated transcripts identified as differentially regulated in Fn CFS were affected. We found that expression of *cidA*, encoding a holin-like protein involved in programmed cell death and extracellular DNA release during biofilm formation, was approximately ten-fold higher (p<0.0001) in the presence of propionate, but was relatively unaffected by acetate or butyrate (Fig. 3B)^28^. Conversely, there was no effect of SCFAs on the expression of *lrgA*, which encodes a putative anti-holin that is antagonistic to CidA^28^. This suggests that decreased *lrgA* expression detected in Fn CFS is likely indepdendent of the SCFAs tested here (Fig. 3C). Despite increased *cidA* expression, SCFA supplementation of LB had marginal effects on biofilm production, with acetate and propionate leading inducing modest but insignificant increases relative to LB alone, and butyrate having no detectable effect (Fig. 3D). The lack of downregulation of *lrgA* under these conditions suggests that sufficient LrgA protein may be available to offset any increased CidA activity. Alternatively, other environmental cues may be needed to enhance biofilm formation under these conditions.

### The *fadX* gene mediates propionate resistance

The most highly induced transcript in *S. aureus* grown in Fn CFS was *fadX*, encoding a putative propionate CoA-transferase, the first in a five gene operon predicted to be involved in fatty acid degradation. Given their annotation, we hypothesized that the *fad* operon may encode a component of the *S. aureus* SCFA stress response. The *fad* genes were induced after growth in Fn CFS and in the presence of propionate and butyrate; their status as an operon was confirmed by obtaining amplicons from cDNA using PCR primer sets that spanned each intergenic region (Fig. 4, Fig. S2). We tested a *fadX* transposon mutant (obtained from the Nebraska Transposon Mutant Library) and its parental strain (JE2) for the ability to grow in 100 mM propionate, and found that *fadX*::tn had a significant growth defect in propionate relative to the wild type (Fig. S3). The mutant grew as well as the parent strain in LB alone, indicating that growth inhibition was specific to propionate in the medium. We then performed dose-response growth curves in six concentrations of sodium propionate, ranging from 100 mM to 3.125 mM in two-fold reductions, and found clear growth differences between JE2 and *fadX*::tn, with the mutant having a defect in media with as low as 12.5 mM (Fig. S4A).

**Figure 4.**
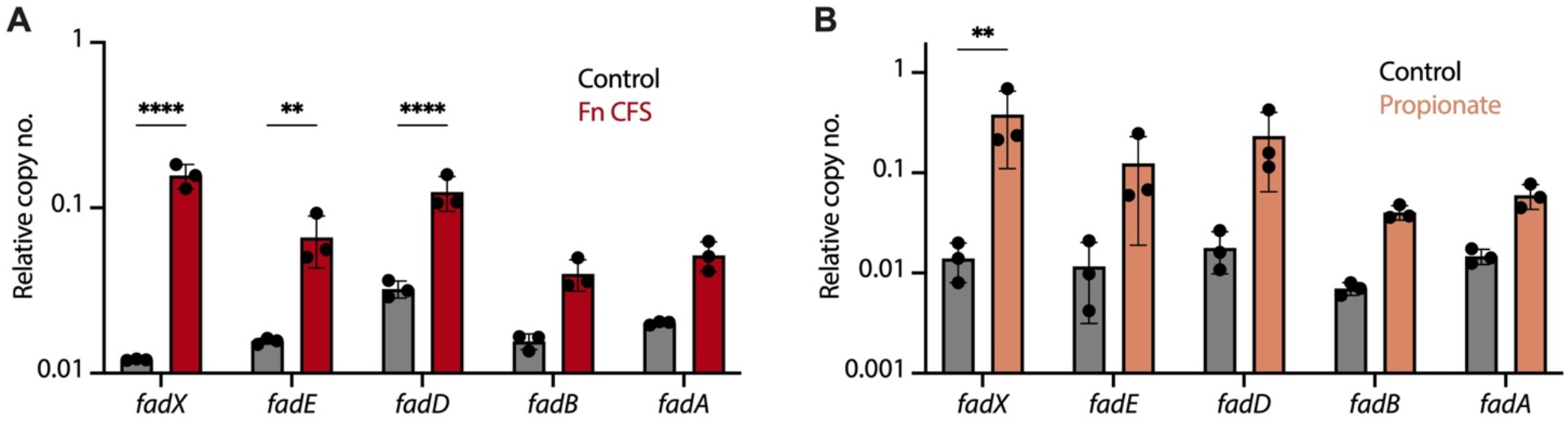
The *fad* operon is induced by propionate and butyrate. **A)** Quantitative reverse transcription PCR was used to detect *fad* operon expression in control (BB) or Fn CFS, or **B)** in LB supplemented with 100 mM sodium propionate. Cultures were grown to an OD_600_ of approximately 0.2-0.3 prior to RNA extraction. Data shown are mean +/− standard deviation of three biological replicates.

To determine if the transposon insertion in the *fadX*::tn mutant disrupted the entire *fad* operon, we performed qRT-PCR and confirmed that the three genes downstream of *fadX* had considerably reduced expression in LB (Fig. S4B). We therefore constructed a ∆*fadX* deletion mutant in the USA300 LAC background and tested its growth in LB supplemented with each SCFA (Fig. 5). Relative to wild type, there was no growth defect in acetate, however there was modest inhibition of the mutant in propionate, with growth curves diverging after approximately 8-10 hours and remaining consistent through the end of the experiments. Neither strain grew well in butyrate, though sporadic growth was detected after ~15 h, irrespective of genotype and only in butyrate. Together, these data implicate FadX in ameliorating or resisting propionate stress, though the mechanism remains unclear. Further, the occasional growth of either strain at later time points in butyrate, but not propionate, provides indirect evidence that these SCFAs may have different mechanisms of *S. aureus* growth inhibition.

**Figure 5.**
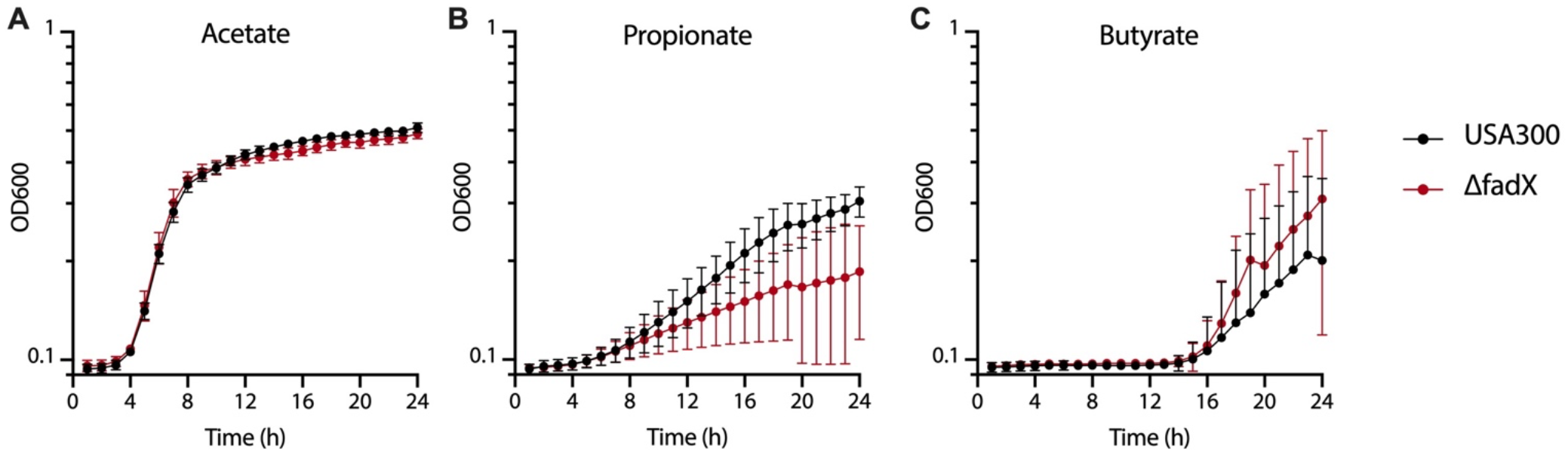
The ∆*fadX* mutant is more susceptible to growth inhibition by propionate than wild type. Combined growth curves (n=4) of wild type USA300 or the ∆*fadX* mutant in 100 mM of the sodium salts of **A)** acetate, **B)** propionate, or **C)** butyrate. Data shown are the mean +/− standard deviation of three biological replicates.

### *codY* mutants are resistant to butyrate

Although occasional growth was detected in LB supplemented with butyrate after ~15 hours of incubation, it consistently occurred in only one of three technical replicates of a given sample (either wild type or ∆*fadX*). These wells were plated onto LB and mannitol salt agar to confirm the absence of contamination and that the observed turbidity was due solely to *S. aureus* growth. To further investigate this phenomenon, we grew the ∆*fadX* mutant for 24 h in LB, then plated ten-fold serial dilutions onto LB agar + 200 mM sodium butyrate. Large colonies were occasionally observed in the 10^-2^ dilutions after overnight incubation, which we assumed to be due to increased resistance to butyrate. Patching these colonies onto the same medium confirmed their resistance phenotype. We interpret these data to mean that butyrate resistant mutants arose spontaneously in LB starter cultures, and that the occasional turbidity in LB + butyrate cultures after ~15 hours represents growth after an extended lag phase resulting from their extremely low starting abundance.

Growth curves in each SCFA were then performed to determine if the large colonies had a growth advantage over the parental strain. We found that all four large colonies grew significantly faster in the presence of butyrate than the parental strain, yet there were no differences in media supplemented with acetate or propionate (Fig. 6A-C). We then sequenced their genomes to identify genetic determinant(s) of butyrate resistance, and found two independent mutations in the gene encoding the GTP- and branched chain amino acid-sensing global regulator CodY. The first mutation resulted in a premature stop codon truncating the protein after 65 amino acids, while the second led to a 20 amino acid deletion from a conserved region of the protein at codons 171-190 (Fig. 6D). We repeated growth curves using a JE2 *codY*::tn mutant (with an intact *fadX* gene) and confirmed that *codY* mutation alone was sufficient to rescue growth in the presence of butyrate (Fig. 7A). Finally, we performed qRT-PCR on JE2 and the *codY*::tn mutant to assay for *fad* operon expression and found that while the operon was modestly induced in the mutant, none of the genes reached significance (Fig. 7B). These data, coupled with the fact that *codY* mutants have similar levels of growth impairment in propionate is further evidence that propionate and butyrate may act on different targets to inhibit *S. aureus* growth.

**Figure 6.**
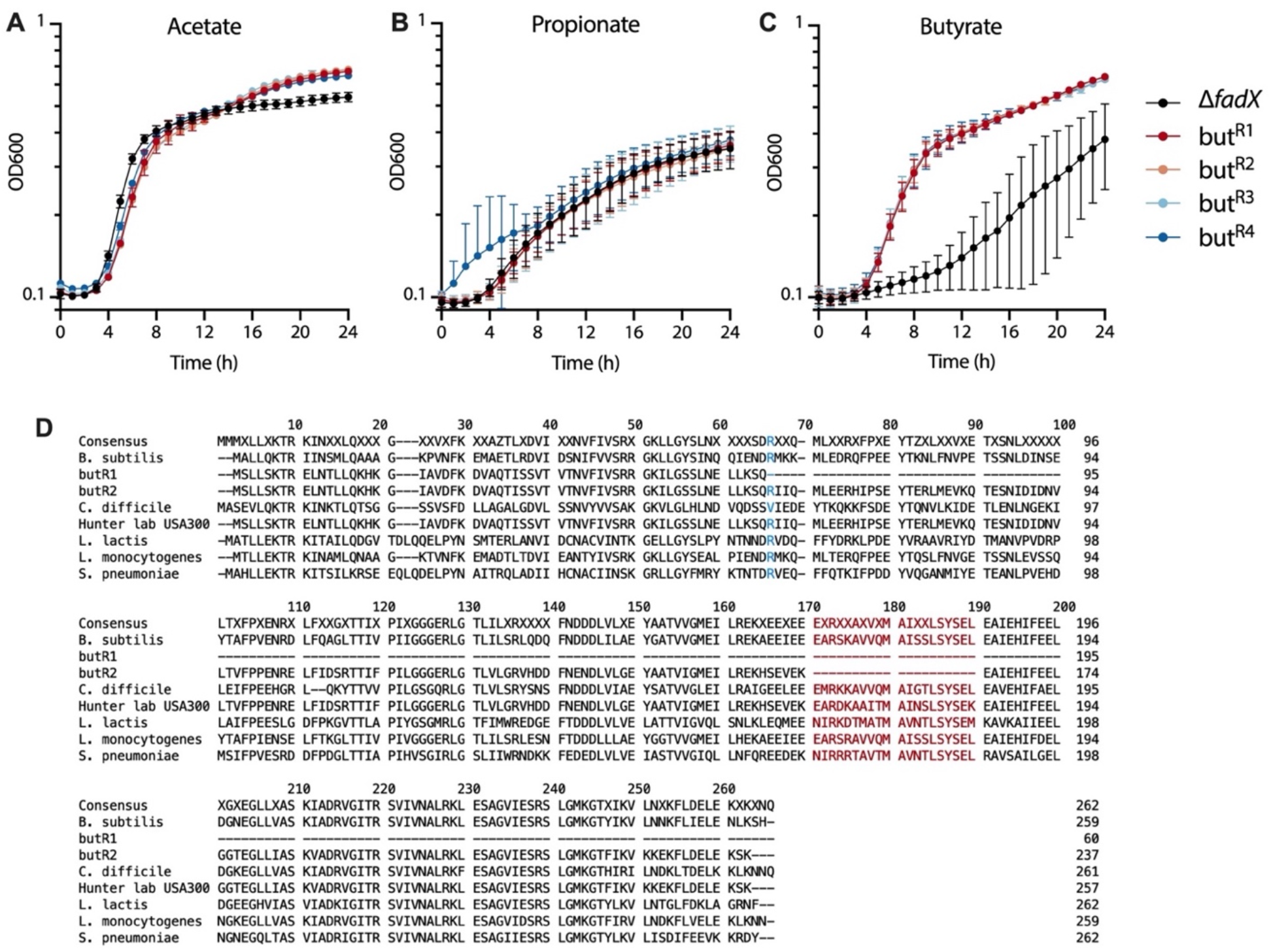
Spontaneous *S. aureus codY* mutants are not inhibited by butyrate. Combined growth curves of the ∆*fadX* mutant and butyrate-resistant derivatives in 100 mM **A)** sodium acetate, **B)** sodium propionate, or **C)** sodium butyrate. **D)** Alignment of CodY protein sequences from diverse Gram-positive bacteria and *S. aureus*, including butyrate resistant mutants (but^R1^ and but^R2^). but^R1^ encodes a premature stop codon at position 66 (blue), while but^R2-4^ mutants have a 20 amino acid deletion from positions 171-190 (red). but^R3^ and but^R4^ mutants were omitted from the alignment as their codY mutations are identical to but^R2^.

**Figure 7.**
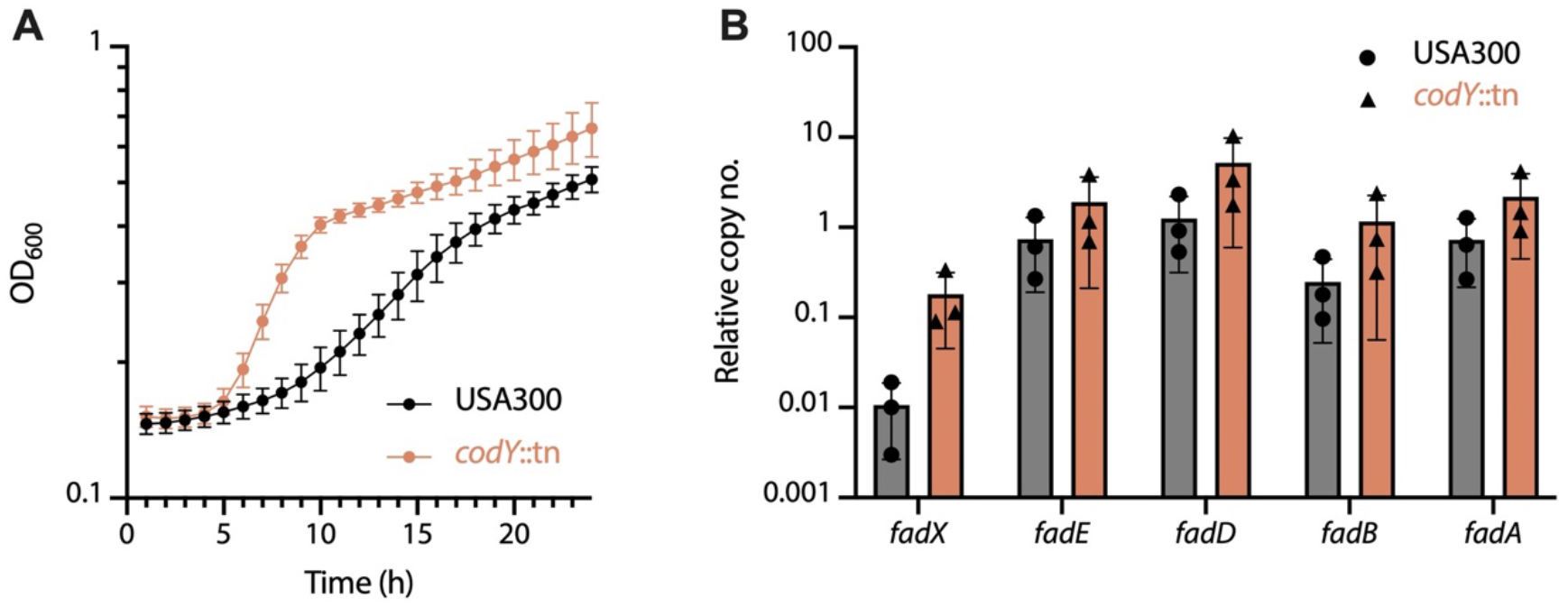
*codY* mutation is sufficient to escape butyrate growth inhibition, though not likely through Fad activity. **A)** Growth of wild type *S. aureus* and a *codY*::tn mutant in 100 mM sodium butyrate. **B)** Expression of the fad operon from the same strains as in A, grown in LB to an OD_600_ of approximately 0.2-0.3.

## DISCUSSION

Although not lethal, CRS remains a significant source of morbidity for a large percentage of the population (~15%), and recalcitrance to antibiotic therapy often requires invasive surgical intervention^5^. The recent advances in bacterial community profiling by 16S amplicon sequencing has revealed extensive colonization of CRS sinus mucus with oral anaerobes and other taxa not frequently observed via traditional culture-based methods^4,6^. While *S. aureus* is appreciated as a significant CRS pathogen, it coexists with these communities and must adapt to the sinonasal microenvironment shaped by both host and microbial processes. Many of the anaerobes associated with CRS release nutrients in the form sialic acid and other carbohydrates, peptides and amino acids, and byproducts of mixed-acid fermentation^6^. One such class of metabolites, short-chain fatty acids, are derived from amino acid-fermenting anaerobes, particularly members of the *Fusobacterium* genus^29^. Our goals in this work were to (i) determine if *Fusobacterium nucleatum*, a model member of the Fusobacteria, could impair *S. aureus* growth, (ii) characterize the response of *S. aureus* to individual SCFAs, and (iii) identify potential mechanisms of SCFA stress.

We show that *F. nucleatum* produces millimolar amounts of the SCFAs acetate, propionate, and butyrate, and that *S. aureus* has an extended lag phase and significant alterations to its transcriptome when grown in *F. nucleatum* supernatants or control medium supplemented with SCFAs. Consistent with a prior study, propionate and butyrate were both inhibitory to *S. aureus* growth, though we found that butyrate was more potent in this regard^26^. Both SCFAs were sufficient to reduce expression of the master regulator of virulence, *agrA*, and alter expression of metabolic pathways (*cidA* and the *fad* operon), in support of the hypothesis that SCFAs were responsible for altered gene expression and the delayed lag phase in *F. nucleatum* supernatants. Reduced *agr*-regulated virulence factor output may alter the inflammatory tone of the CRS sinus environment, with the host response instead being directed towards members of the anaerobic community rather than *S. aureus*. In support of this idea, CRS patients have been reported to have circulating antibodies targeting *Fusobacterium* and *Prevotella*, members of which were enriched in CRS sinus mucus in our previous study, and that they show a decline in these antibodies after successful antibiotic therapy^6,30^. Alternatively, butyrate produced by anaerobes in the CRS sinus environment may occasionally select for *S. aureus codY* mutants, whose growth inhibition would be relieved^31^. Such mutants overproduce numerous virulence factors, and as such, may exacerbate the inflammatory response^32^. Whatever the case, the recent development of robust animal models of CRS will facilitate our ability to test these hypotheses *in vivo*^33,34^.

While the mechanisms of action of propionate and butyrate on *S. aureus* are still unclear, our data suggest that they may act on different targets. Propionate induced expression of the *fad* operon and a *fadX* mutant exhibited worse growth in its presence than did the wild type strain. The *fad* operon annotation suggests a role in fatty acid degradation, though it is unlikely that it acts directly on propionate, as *S. aureus* showed no evidence of metabolizing it over time (Fig. 1C). Another possibility is that propionate induces lipid membrane stress, and the Fad proteins may act to degrade or repair damaged lipid species. This consistent with recent findings from human gut commensal Bacteroides, where butyrate (rather than propionate) induced membrane stress in a species- and context-dependent manner^35^. Butyryl-CoA levels were increased by Acyl-CoA enzymatic activity, suggesting that other CoA-regulated enzymes could be starved of an essential cofactor, likely imparing several metabolic processes. While butyrate also induced expression of *fadX*, we did not detect a mutant phenotype in LB supplemented with 100 mM butyrate, suggesting that the other members of the *fad* operon may compensate for the loss of *fadX* under these conditions, or that they are less important for the response to butyrate. Interestingly, we readily obtained spontaneous butyrate resistance in the form of *codY* mutations, while we did not for propionate. Further, butyrate resistant *codY* mutants were as sensitive to propionate as the parent strain, suggesting that each SCFA may have unique mechanisms of action on *S. aureus*. CodY is an allosteric transcriptional regulator whose affinity for specific motifs in promoter regions is dictated by the levels of GTP and branched chain amino acids in the cell^36^. As it regulates hundreds of genes in *S. aureus*, determining which CodY target(s) are responsible for bypassing butyrate-mediated growth inhibition are beyond the scope of this work, though transposon screens in a *codY* mutant background may prove fruitful in this regard.

In summary, we have identified a possible mechanism by which anaerobic bacteria in polymicrobial airway infections may influence *S. aureus* growth and physiology via the activity of the short-chain fatty acids propionate and butyrate, and have identified the *fad* operon and the CodY regulon as possible mechanisms of resistance, respectively. Our study has some limitations, as the experiments were performed in *F. nucleatum* supernatants *in vitro* and in defined media conditions rather than in the context of intact anaerobic communities or two-species co-cultures. Additionally, our experiments lack the potential contributions of the host, such as reactive oxygen and nitrogen species, or cationic antimicrobial peptides^37,38^. Despite these limitations, the genetic approach taken here is informative and amenable to translation into animal models for further dissection of the effects of CRS bacterial communities on *S. aureus* pathogenesis.

## ACKNOWLEDGEMENTS

We acknowledge the UMN Genomics Center and Paige Marsolek for NanoString assistance, and members of the Hunter laboratory for critical feedback on the manuscript. This work was supported by a National Institute of Dental and Craniofacial T32 Fellowship (#T90DE0227232) awarded to JRF, a National Heart, Lung, and Blood Institute Research Project Grant (1R01HL136919) to RCH, and an Administrative Research Supplement (HL136919-03S1) to ARV. The funders had no role in study design, data collection and interpretation, or the decision to submit the work for publication.

## Notes

### Competing Interest Statement

The authors have declared no competing interest.

