## Supplemental Tables S1, S2, Figures S1-S4 for "*Staphylococcus aureus* overcomes anaerobe-derived short-chain fatty acid stress via FadX and the CodY regulon"

**Fletcher et al.**

### Supplemental Data

**Table S1. Bacterial strains and plasmids**

| Strain or Plasmid | Characteristic | Source |
|---|---|---|
| ***S. aureus*** | | |
| USA300 | Strain LAC | |
| USA300 Δ*fadX* | USA300 with a deletion of *fadX* | This study |
| JE2 | USA300 JE2 | Fey et al., 2013 |
| JE2 *fadX*::tn | JE2 with a *bursa borealis* insertion in *fadX* | Fey et al., 2013 |
| JE2 *codY*::tn | JE2 with a *bursa aurealis* insertion in *codY* | Fey et al., 2013 |
| **Plasmids** | | |
| pIMAY | mutagenesis | Monk et al., 2012 |
| pAH1 | P*agr-mCherry* | Malone et al., 2009 |
| pCM29 | P*sarAP1-sGFP* | Pang et al., 2010 |
| p*fadX*c | pCM29 with GFP replaced by *fadX* coding sequence | This study |

**Table S2. Primer sequences**

5'fadX.up.KpnI - TTATAGGTACCGTTGATTGTGTACTTGCTGATGAC
3'fadX.up.SOE - TTAGCTTCACTTTCACCATGTGATTTGTTTCAAGCAAGTCACCTC

5'fadX.down.SOE - AAACAAATCACATGGTGAAAGTGAAGCTAAAGGGGGTATACTAATG
3'fadX.down.SacI - TTATAGAGCTCCACCTTTAGGTGATCCGGTTG

5'fadXc_KpnI - TTATAGGTACCGTAAGGGTTTACACAAAGTGTAAAAACGC
3'fadXc.EcoRI - TTATAGAATTCCCTTTAGCTTCACTTTCATACTTTATGAATTGATTG

5'fadXE.junction - ATGGCTTTTAAACCAATTATTGCTG
3'fadXE.junction - GATCAATAACGGCAGGCTTGTC

5'fadED.junction - TGTACCAGTGACACATATGCCG
3'fadED.junction - CGACTTCACCGTCTGTTAAACC

5'fadDB.junction - AATGCCTTAGTAATTGGACGCG
3'fadDB.junction - CAAAAAGTGCTGCCAGTTGAGC

5'fadBA.junction - CGCCACAATTTTAGCGGGTG
3'fadBA.junction - GTCTTTCGTGGAATAATGCGCC

5'fadX.q - TTGATGTGGCACTACTGAGAGG
3'fadX.q - GATTGGCTTTCGCGTTTAATGC

5'fadE.q.Sa - TGGTGACTTAGCGAAGATGGAC
3'fadE.Sa.q - GGTCTACTAGTGGATGCTCAGC

5'fadD.q.Sa - AGTCAGACCAGAACAAGATGGC
3'fadD.q.Sa - CGCCTGCTCTCGTTGAATAAAG

5'fadB.q.Sa - GTGGATGCTTTAACTGGGCAAG
3'fadB.q.Sa - CTTCAGGTACTTGTTGCATGCC

5'fadA.q.Sa - CTGGTGGCGTTGAATTGATGAG
3'fadA.q.Sa - AACCCATAGGATATGACGCACC

5'cidA.q - ACCGCTAACTTGGGTAGAAGAC
3'cidA.q - AGCGTAATTTCGGAAGCAACATC

5'lrgA.q - GGTGCTGTTAAGTTAGGCGAAG
3'lrgA.q - AATGGTGCTTGGCTAATGACAC

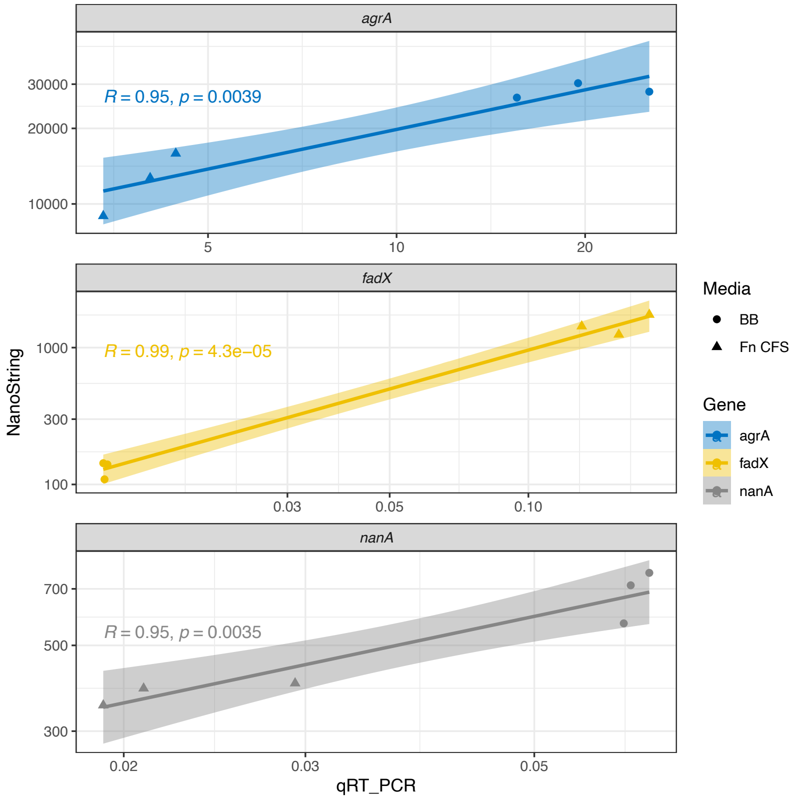


**Figure S1. Validation of select NanoString results (y-axis) by quantitative reverse transcription PCR (x-axis).** Complementary DNA was generated from the RNA used in the NanoString assay, then subsequently used as template for qRT-PCR assays. Each qRT-PCR dataset was highly correlated with its corresponding NanoString result.

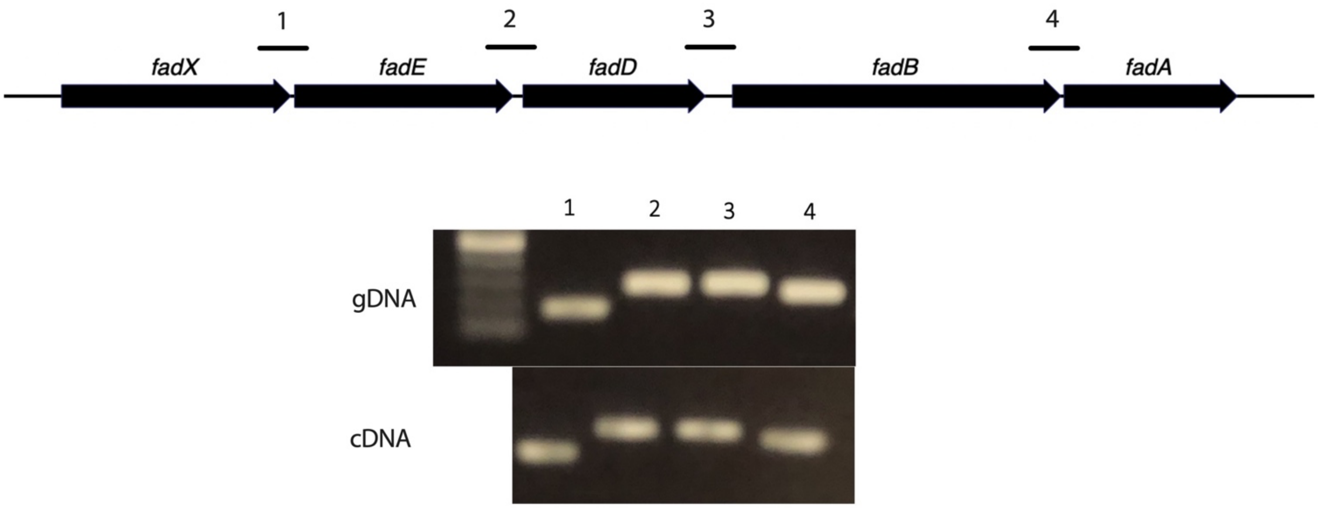


**Figure S2. *fad* genes are operonic.** PCR primers targeting the numbered intergenic regions between each *fad* gene were used to produce amplicons from genomic DNA and cDNA.

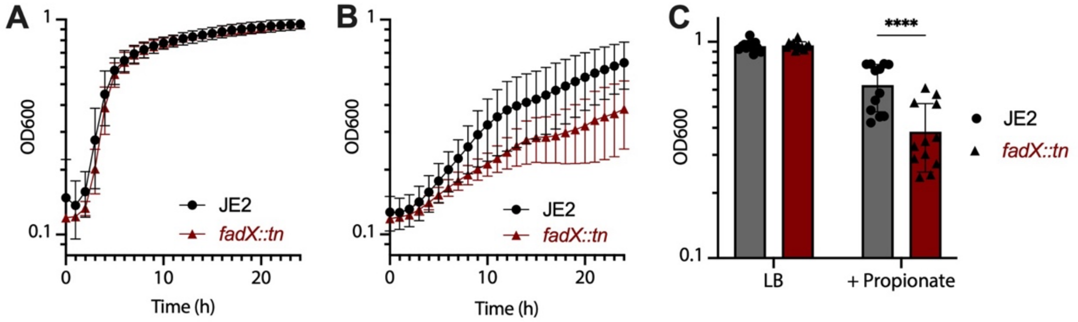


**Figure S3. The *fadX*::tn mutant is significantly more sensitive to propionate compared to the wild type. A)** Growth of the parent strain *S. aureus* JE2 and *fadX*::tn in LB, or **B)** LB supplemented with 100 mM sodium propionate. **C)** $OD_{600}$ values of each strain (n=4 biological replicates) at 24 h in LB or LB + propionate.

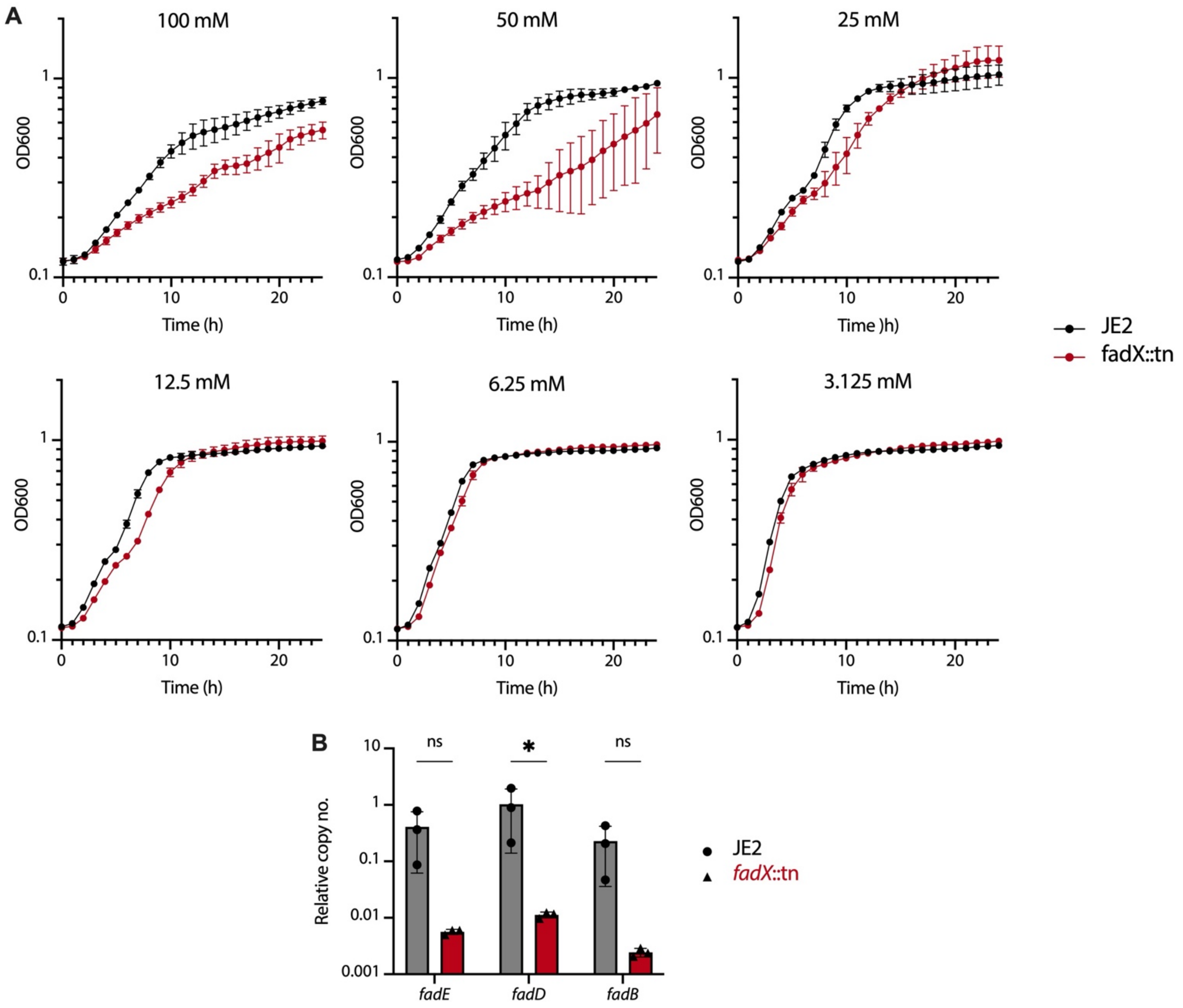


**Figure S4. *fadX* is required for optimal growth in physiologically relevant levels of propionate. A)** Growth of *S. aureus* JE2 and *fadX*::tn in LB supplemented with various concentrations of sodium propionate. **B)** qRT-PCR data showing that the three genes immediate downstream of *fadX* have reduced expression in the *fadX*::tn mutant relative to wild type JE2, indicating polar effects of the transposon insertion.
